# A social-ecological geography of southern Canadian Lakes

**DOI:** 10.1101/2023.03.09.531893

**Authors:** Andréanne Dupont, Morgan Botrel, Nicolas Fortin St-Gelais, Timothée Poisot, Roxane Maranger

## Abstract

Anthropogenic pressures, including urban and agricultural expansion, can negatively influence a lake’s capacity to provide aquatic ecosystem services (ES). However, identifying lakes most at risk of losing their ES requires integrating information on lake ecological state, global change threats, and ES demand. Here, we provide a social-ecological framework that combines these features within a regional context based on an ecological evaluation of the state of 659 lakes across Canada. From deviation of impacted lakes to reference ones, we identified much higher concentrations of total nitrogen and chloride as the main indicators of altered lake ecological state in all regions identified. Lake ecological state was mapped using an additive colour model along with regional scores of threat levels and recreational ES demand. Population density and agriculture were linked to high lake vulnerability. Lakes in Southern Ontario were most concerning, being highly altered, under threat, and heavily used. Lakes near urban centers along coasts were altered and used, but less threatened, whereas those in the Prairies were altered and threatened, but less used. Our novel framework provides the first social-ecological geography of Canadian lakes, and, is a promising tool to assess lake state and vulnerability at scales relevant for management.

**Plan language summary:** Plain language title: Assessing overall lake health across Canada to identify sites for restoration and conservation

Canadians love to swim, fish, and navigate in and on the countless lakes across the country. But Canadian lakes are under a considerable amount of pressure from human activities in their watershed. The expansion of cities, intensive farming, wetland loss, and industrial development all results in the transfer of pollutants to aquatic habitats, threatening the health of lakes and the ecosystem services they provide. Where are lakes being used across Canada? What condition are they in and is their use under threat from different pressures? To answer these questions, we combined information from many different sources, including a national scale lake assessment, through the NSERC Strategic Network Cluster Lake Pulse to create the first social-ecological geography of southern Canadian lakes. Regionally specific baseline conditions were established from lakes considered healthy due to limited human activities in their watershed. When lakes with impacted watershed were compared to healthy ones within their specific region, two early warning signals of human pressure, pollution from nitrogen found in fertilizers and sewage, and chloride found in road salt, determined whether a lake was altered. We combined these two health indicators, with information on future potential lake threats and use by the population for recreational purposes. Using a colour-coded mapping technique, we were able to identify regions where lakes were altered, threatened, and used. These regions occurred primarily around dense urban areas, of southern Ontario and Quebec, and major cities on the east and west coast. Lakes were altered and threatened, but seemingly less used in the Prairie Provinces. The novel approach is very adaptable, easy to understand, and can be used at more regional levels for management to determine priority sites for conservation and restoration, as well as in science communication to describe overall lake health.

## Introduction

With over 900,000 lakes shaping its landscape, Canada is the most lake-rich country in the world (Minns et al., 2008). These lakes hold inherent cultural, economic, and ecological value to the peoples of Canada, including First Peoples, and provide a wide variety of aquatic ecosystem services (ES), ranging from drinking water to recreational opportunities such as fishing and swimming, which all contribute to human well-being (Bradford et al., 2017; Keeler et al., 2012; Millennium Ecosystem Assessment, 2005). In fact, over 90% of Canadians rely on lakes and rivers for their drinking water (Environment Canada, 2011), and recreational fisheries represent a $2.5 billion-dollar (CAN) industry (Fisheries and Oceans Canada, 2019). However, these valuable ecosystem services are under threat by anthropogenic pressures that alter surface water quality (Brauman et al., 2007; Dugan et al., 2017; Keeler et al., 2012). These include local land use changes, increasing nutrient and pollutant inputs from the watershed as well as regional climate change that can severely disrupt a lake’s hydrological regime, thermal structure and internal processing, thus altering a lake’s chemical and biological properties (Carpenter et al., 2011; Jeppesen et al., 2014; Schindler, 2001; Vincent, 2009). Human dependence on aquatic ES is also expected to increase in the face of climate adaptation and increasing population growth (Cimon-Morin et al., 2014; Guo et al., 2010) where water use and withdrawal will need to be adjusted for changes in availability or quality given demands (Bakker & Cook, 2011; Brandes & Ferguson, 2004; Schindler & Donahue, 2006). Therefore, there is an urgent need to protect lakes and the ES they deliver (Baron et al., 2002; Culhane et al., 2019; Hossu, 2019), which begins with understanding the regional context in which a lake’s ES are threatened.

The response of a lake to different anthropogenic pressures, be it changes in climate or land use is highly variable given the diversity of lake and watershed traits. A lake’s morphometry, regional geology, watershed topography, soil composition, and land cover are all factors that can either amplify or mitigate its response to various pressures (Jeppesen et al., 2009; Nõges, 2009; Tang et al., 2005). Combined, these biophysical features will influence a lake’s baseline conditions and resilience in providing different aquatic ES. For example, human activities in watersheds increase nutrient loads to lakes (Nielsen et al., 2012), but lakes with a high drainage ratio (watershed area: lake area) tend to be naturally more enriched, receiving greater terrestrial inputs and as such may be more susceptible to eutrophication (Knoll et al., 2015). Likewise, lakes from the prairies are geologically embedded in nutrient-rich sedimentary rocks and have naturally higher phosphorus (P) concentrations as compared to those in the Boreal Shield, for example (D’Arcy & Carignan, 1997; Prepas et al. 2001), resulting in different reference conditions of unimpacted ecosystem. Given the broad range in topographies and surface geologies across Canada (Boyce, 2015), the limnological landscape is likely highly diverse. Therefore, a lake’s state needs to be considered within a regional biophysical context (Kumagai & Vincent, 2003; Soranno et al., 2015) to establish how severely it deviates from its natural state to implement efficient lake management strategies and set reasonable restoration targets.

Beyond understanding how a lake deviates from a regional reference in terms of ecological state, protecting a lake’s capacity to provide ES into the future, requires regional social information on anthropogenically driven threats to that state, and on ES use. To our knowledge, this type of integrative assessment does not exist, and one of the challenges of creating it at a national scale, is that the required information is often fragmented across various jurisdictions. For example, although there are many water quality monitoring programs across Canada, these initiatives are either carried out at provincial or more local levels, and methods used to compare lakes are not standardized (Huot et al., 2019). In terms of threats to lakes, there is a high degree of regional and social variability in land use change or impacts from invasive species (Mandrak & Cudmore, 2010; Minns et al., 2008; Walsh et al., 2016), and although climate change may be more ubiquitous, climate effects vary spatially across Canada (Barrow et al. 2004; Murray et al. 2023). Which ES are valued and how intensively lakes are used for those services will also differ across regions. This may be related to the proximity of human populations to lakes, but may also differ as to which ES may be available as a function of lake type (Hunt & Dyck, 2011). Regardless, knowing whether a lake’s ES is at risk or not, requires an understanding of whether the ES is in demand (Chaplin-Kramer et al., 2019; Mitchell et al., 2021; Villamagna et al., 2013), and this will vary as a function of not only biophysical conditions, but social interests.

Given the importance of understanding how lakes with different biophysical traits respond to change, the overarching goal of this study is to generate the first social-ecological geography of Canadian lakes. With the purpose of sustaining water quality and aquatic ES into the future, this study provides a regionally based evaluation of lake ecological state by first determining the reference biophysical and chemical properties of unimpacted lakes, and estimating the deviation of impacted lakes from this regional reference. An integrated regional spatial analysis of lake ecological state, cultural ES demand, and impending threats from different anthropogenic pressures using a colour additive model creates a novel and easily interpreted social-ecological portrait of lakes across the country. We suggest that the integration of these three components provides a holistic approach to assess overall lake ecosystem state or health, and enables the identification of the most vulnerable lakes within a region for targeted protection of the ES valued by the local population.

## Methods

### Study area and Datasets

To assess the three components of the socio-ecological geography, data were collected or retrieved from three large-scale Canadian datasets: (1) lake biophysical and chemical characteristics, from NSERC Canadian Lake Pulse Network – Lake Survey (Huot et al., 2019), (2) threat level, from World Wildlife Fund (WWF) Canada – Watershed Reports (2017) and (3) recreational uses as ES, from Statistics Canada – Survey on the Importance of Nature to Canadians (Special Surveys Division, 1996).

#### Lake biophysical and chemical characteristics

##### Lake selection and classification

One of the goals of the NSERC Canadian Lake Pulse Network was to conduct a national assessment of lake health (Huot et al., 2019). Description of Canada’s physiographic regions as well as information on land use and population density are available in supplementary information (supplementary material Text S1, Figures S1a and S1b). A total of 664 lakes in 12 Canadian ecozones were sampled by Lake Pulse over three yearly summer sampling campaigns (2017 – 2019). Lakes were selected based on road accessibility and three criteria in a stratified random design: ecozone, lake size, and human impact index (Huot et al., 2019). Ecozones are ecological regions delineated based on similar climate and geological characteristics such as dominant vegetation and landform features (Ecological Stratification Working Group, 1996). Most lakes were sampled in the 9 southernmost ecozones (Boreal Shield, Atlantic Maritime, Boreal Plains, Prairies, Montane Cordillera, Pacific Maritime, Boreal Cordillera, Taiga Cordillera, Atlantic Highlands, and Semi-Arid Plateaus) with a few additional lakes sampled in 3 other ecozones located further north (Taiga Plains, Boreal Cordillera, and Taiga Cordillera). As such this assessment is largely restricted to southern areas of Canada, nearer to urban areas and major highways. Lakes were also divided in three size classes: small (0.1 – 0.5 km^2^), medium (0.5 – 5 km^2^) and large (5 –100 km^2^) and in three human impact classes (low, moderate, high). Impact classes were defined as a function of the proportion of land use and land cover (LULC) in each watershed where urban areas, mines, and agriculture had the highest human impact value (1), pasture and forestry had a medium impact value (0.5) and natural landscape had no impact (0; Huot et al., 2019).

##### Lake sampling and laboratory analysis

Lakes were sampled between July and the beginning of September using standardized protocols (NSERC Canadian Lake Pulse Network, 2021) with approximately 100 variables collected in each lake. For this study, we selected 20 core variables that are relevant for determining the biophysical and ecological state of lakes in Canada, and removed those that were capturing redundant information (*r* > 0.8, Table 1). We grouped variables in those that describe physical features such as lake basin and morphometry, and those that describe water chemistry (including chlorophyll *a* – Chl a - as a biological indicator). For chemical variables, we chose variables that reflected lake productivity (TN, TP, Chl a, secchi depth), geology (chloride, pH and conductivity), and carbon inputs (DOC, coloured dissolved organic matter – CDOM – absorption coefficients of CDOM at 254 and 440 nm wavelength were specifically chosen: abs440 and SUVA254). All variables considered for this study were sampled from the epilimnion at the deepest sites of the lake (based on a bathymetric map or a rapid depth survey by sampling teams) from a boat.

**Table 1.** Selected variables and their units.

| Type of variable | Variable | Unit | Transformation |
| --- | --- | --- | --- |
| Chemical | Total nitrogen (TN) | mg L<sup>-1</sup> | Log |
|  | Total phosphorus (TP) | µg L<sup>-1</sup> | Log |
|  | Chlorophyll *a* (Chl *a*) | µg L<sup>-1</sup> | Log |
|  | Dissolved organic carbon (DOC) | mg L<sup>-1</sup> | Log |
|  | Secchi | m | Square root |
|  | Abs440 | 1/m | Log |
|  | SUVA254 | L/mg-m | Square root |
|  | Chloride (Cl<sup>-</sup>) | mg L<sup>-1</sup> | Log |
|  | Sulfate | mg L<sup>-1</sup> | Log |
|  | C:N ratio | - | Log |
|  | C:P ratio | - | Log |
|  | N:P ratio | - | Log |
|  | pH | - | - |
|  | Conductivity (specific) | µS/cm | Log |
| Morphometric | Lake area | km<sup>2</sup> | Log |
|  | Mean depth | m | Log |
|  | Mean slope | - | Log |
|  | Altitude | m | Square root |
|  | Drainage ratio | - | Log |
|  | Dynamic ratio | - | Log |
| LULC | Fraction of land-use categories in the watershed (Agriculture, forestry, mines, pasture, urban, water or natural landscape) | % | - |

Measurements of secchi depth (secchi disk), pH (RBR multi-parameter sonde or a Hannah pH-meter), and conductivity (RBR multi-parameter sonde) were taken on site. For all other chemical variables, water samples were collected in the field with a tube sampler, placed in pre-cleaned conditioned carboys on the boat, transferred into individual samples on shore, stored, and analyzed later. TN, fixed with H_2_SO_4_ and stored at 4°C, was analyzed with a continuous flow analyzer (OI Analytical Flow Solution 3100) using the alkaline persulfate digestion method, coupled with a cadmium reactor (Patton & Kryskalla, 2003). TP, stored at 4°C, was analyzed by colourimetric detection with a spectrophotometer (*Ultrospec 2100pro*) at 890 nm after potassium persulfate digestion (Wetzel & Likens, 2000). Chl a samples were filtered in the dark on GFF filters, and stored at -80°C until measured by fluorometry using a Turner Trilogy Laboratory Fluorometer (MacIntyre & Cullen, 2005; Welschmeyer, 1994). DOC, prefiltered using a 0.45 μm syringe filter, fixed with HNO_3_ and stored at 4°C, was analyzed with an OI Analytical Aurora 1030W TOC Analyzer following persulfate oxidation. For CDOM absorption coefficients, samples were prefiltered at 0.45 μm using a syringe and stored in an amber bottle at 4°C, and measured by spectrophotometer after a final filtration at 0.22 μm (Brezonik et al., 2019). We calculated the specific ultraviolet absorbance at 254 nm (SUVA254) by dividing the absorbance at 254 nm (1/m) by the DOC concentration (mg/L) (Weishaar et al., 2003) while the absorbance at 440 nm (abs440) was used as is. Chloride (Cl^-^) and sulfate (SO_4_ ) were prefiltered using a 0.45 μm syringe filter, fixed with HNO_3_, stored at -20°C, and analyzed by ion chromatography using a Dionex DX-600 Ion Chromatography following the US Environmental Protection Agency Anion - Method 300.1 (US EPA, 1997). We calculated Carbon (C), Nitrogen (N) and Phosphorus (P) ratios from the molar concentrations of DOC, TN and TP.

#### Physical variables and land use and land cover

Physical variables included lake depth, size, and shape along with watershed topographic features. Lake area was obtained from the National Hydro Network (Natural Resources Canada, 2015). Mean depth was retrieved from *lakemorpho* 1.2.0 package in R (Hollister & Stachelek, 2017), which estimates the average depth of lake by dividing lake volume by lake surface area. Mean slope is the mean of percent rise inside the watershed calculated from the difference in altitude obtained from the Canadian Digital Elevation Model (CDEM) data layer (Natural Resources Canada, 2013). Drainage ratio, an indicator of the amount of external loading to a lake, was calculated as watershed area divided by lake area (Kalff, 2002), and the dynamic ratio, an indicator of the proportion of the lake bottom that is subject to resuspension (low value bowl-like and high value dish-like lakes), by dividing the square root of lake area by the mean depth (Håkanson, 1982). Land use and land cover (LULC) was derived from geographic information system (GIS) and calculated as the fraction of the lake watershed area covered by agriculture, forestry, mines, pastures, urban areas, water, or natural landscapes (see Huot et al., 2019 for more information).

#### Determining lake ecological state

All statistical analyses were performed in R v. 4.0.2 (R Core Team, 2020). Five lakes were considered outliers and removed from the dataset: Kississing Lake which exceeded the size range with an area of 266 km^2^, Embrees Pond which was very saline and likely connected to the sea, and the 3 lakes in the Taiga Cordillera because of lack of representation in this ecozone. Our final dataset therefore contained 659 lakes from 11 ecozones. Missing data for chemical variables were imputed by a random forest algorithm using the “missForest” package (Stekhoven & Buhlmann, 2012). Most variables had less than 12% of missing data, except for pH at 21%. In addition, and to reduce distributions asymmetry, variables were log or square root transformed (Table 1). To choose the appropriate transformation, we performed Shapiro-Wilk Normality Test using *shapiro.test()* function and visually assessed distributions using histograms.

To determine the natural reference conditions of lakes in different regions across Canada, we used the chemical and morphometric characteristics of lakes that were classified as low human impact within the different ecozones (Huot et al., 2019). These had a median of less than 5% of watershed within a given region (Figure S2).We performed a linear discriminant analysis (LDA, Legendre et Legendre 2012) to determine how these potential reference lakes differed across ecozones following Borcard et al. 2018 with the MASS, vegan and rrcov packages (Venables & Ripley, 2002, Oksanen et al., 2020, Todorov & Filzmoser, 2010). The descriptors (morphometric and chemical variables) were standardized to compute discriminant functions (rather than identification functions) to quantify the relative contribution of the variable to the discrimination between groups (Borcard et al., 2018). Mean depth and nutrient ratios were excluded from this analysis due to collinearity with other variables. For each group, we computed the percentage of lakes correctly classified by the discriminant functions.

To evaluate the difference between impacted and non-impacted lakes within each lake region, we calculated the deviation of impacted lakes from regional reference conditions. In order to multiple variables with different units, standard scores were calculated per lake region for each variable using all lakes (including low, medium, high human impact). The regional lake reference conditions (or the “baseline values”) were estimated for each variable by averaging the standard score of their low human impact lakes. The deviation from reference conditions was calculated as the standard score of high and moderate human impact lakes minus the baseline values. Thus, the baseline values were set to zero for every region and allowed the comparison of deviations among different lake regions.

To visualize the correlation between chemical variables and the more fixed landscape components of lake shape and anthropogenic land use in the watershed, we used a Principal Component Analysis (PCA) with passive projection estimated from the vegan package (Oksanen et al. 2019). The deviation in morphometric and land use variables, which are considered fixed in space, were used to determine the principal components, and the chemical variables were then passively projected onto these principal components. The projected chemical variable represents the linear trend with the environmental variables (morphometry and land use).

#### Threat level

Based on threats to drinking water sources and aquatic ecosystems health identified by Environment Canada (Environment Canada, 2001), WWF’s 2017 Watershed Report (WWF-Canada, 2017) developed seven indicators of the most significant threats currently facing Canada’s watershed, namely pollution, habitat loss, habitat fragmentation, overuse of water, invasive species, flow alteration, and climate change. Among the seven threat indicators reported by the WWF, we selected five that apply to lakes (pollution, climate change, invasive species, overuse of water and habitat loss). Flow alteration and habitat fragmentation were discarded in part because these threats apply more to rivers as well as some of the built-in redundancy of some of these indicator threats. The remaining five indicators were calculated based on more specific sub-indicators that are of growing regional concern: pollution includes point source pollution, pipeline incidents, transportation accidents, and agricultural contamination risks from nitrogen, P, and pesticide release; habitat loss includes land use and land cover as well as forest loss; overuse of water indicates the ratio of water withdrawal to water supply; invasive species indicates the presence of invasive species; and climate change includes summer and winter temperature anomalies and spring and summer precipitation anomalies.

The presence and magnitude of these threats were assessed for Canada’s 25 Pearse watersheds and their 167 sub-watersheds (WWF-Canada, 2015). Data was gathered by the WWF to assess the level of threats for each of the indicators at the sub-watershed level and to establish a relative ranking of threats across the country. Threat categories (ranging from “none” to “very high”) for individual indicators were defined according to a regular interval classification system (WWF-Canada, 2015). For more details on the methods, see WWF’s 2017 Watershed Report “Threat methodology” (https://watershedreports.wwf.ca/#intro). The metric we used for this study is the threat score for each indicator by sub-watershed.

#### Recreational ecosystem service use

Statistics Canada conducted the Survey on the Importance of Nature to Canadians among 87,000 Canadians aged 15 years and older in 1996. This survey assessed the social and economic value that Canadians place on nature-related recreational activities (Environment Canada, 1999). Of interest to our study is the “water and recreation” theme of this survey, which provides information on 4 freshwater recreational uses, considered cultural ES i.e. 1) participation in swimming and beach activities, 2) sport fishing, 3) canoeing, kayaking and sailing, and 4) power boating. The metric we used to assess ES demand in this study is participation frequency (in days per km^2^ of major drainage area) for these 4 recreational water activities. The frequency was calculated by major drainage area (11 in Canada): Maritime Provinces, St. Lawrence, Northern Quebec and Labrador, Southwestern Hudson Bay, Nelson River, Western and Northern Hudson Bay, Great Slave Lake, Pacific, Yukon, Arctic, and Mississippi as defined by the Standard Drainage Area Classification (Natural Resources Canada, 2003). Data on recreational use were adjusted to reflect today’s population in each major drainage area by multiplying the use frequency by the ratio of current population (2016 census) to 1996 population.

### Social-ecological geography: creating gradients across Canada

We overlayed the three components of change in lake ecological state, threat level, and recreational ES to create a socio-ecological geography of Canadian lakes. The map represents a quantitative approach to assess the vulnerability of lakes and the recreational ES they provide to anthropogenic pressures. To produce the map, we used the chemical variables of the impacted lakes that consistently and most strongly deviated from regional baseline values as indicators of alteration of lake ecological state. We assigned a default value of 0 to the low human impact lakes (*n* = 273) since they were used to establish the reference of their region. Given that the selected indicators were typically higher in the impacted lakes compared to the reference, the moderate lakes that had a negative deviation value (*n* = 45) were also set to 0 and considered unaltered. To obtain an overall score for all threats combined, we averaged the scores for the five selected threats within the sub-watershed. The data for the four services were summed to give a total number of days of recreational use per major drainage area.

To combine all three components, each sampled lake was given a score for threat level, lake state and recreational ES use according to its geographical position. Each component was assigned a colour: green for lake state, red for threat level, and blue for recreational use and was scaled from 0 to 255. Zero (0) represents the lowest value in Canada and appears in black, while 255 represents the highest value in Canada and results in the brightest colour (Figure 1a). A red, green, blue (RGB) additive colour model was used to combine all three components on a single map and produce an assortment of colours (Figure 1b). The colour gradient helps to identify areas with similar patterns across Canada. For example, a red area indicates high threat level, a green area indicates a highly altered lake ecological state, and a blue area indicates high recreational ES demand. When all three colours are strongly present, the result is white. This means that the recreational services being used are at risk of being lost because the state of the lake and its threat level are of concern. Other possible combinations of two of these colours will result in the following colour code: when recreational services are being used and the lake is altered, but not threatened cyan; when services are being used and there is a high level of threat, but not altered, magenta; and when the lake is altered, and the threat level is high, but not used, yellow. Ultimately, the lakes to be monitored are located in the magenta, cyan, and white zones because their recreational services are used, and at risk of being lost due to threats (magenta), lake state (cyan), or a combination of both (white).

**Figure 1.**
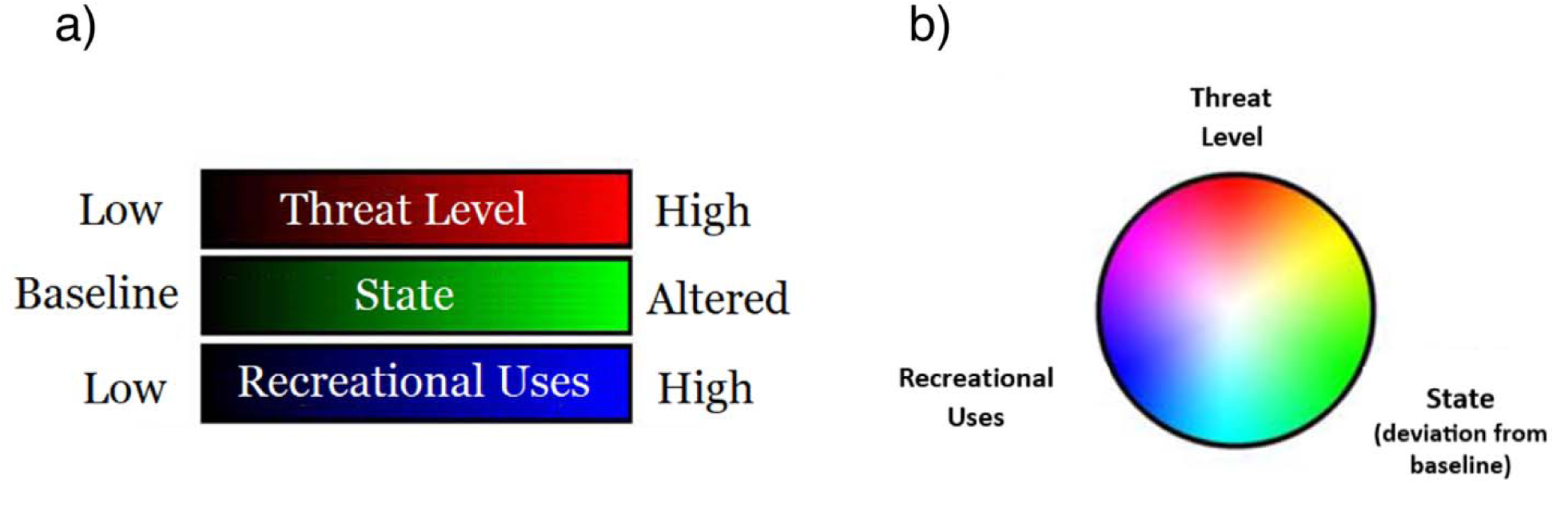
a) Red, green, blue colour gradients for the three components (0 to 255). b) Red, green, blue (RGB) additive colour model (adapted from Seekell et al., 2018) and its resulting gradients.

To generate a continuous map over Canada, sampled lakes polygons were converted to points based on their centroid. Points were interpolated to a continuous raster surface using ordinary spherical kriging with a variable 12 points search radius. The surface of the raster output was then smoothed using a 100 km radius focal mean. Subsequently, a three-band composite image was created by assembling the rasters of all three metrics (i.e., threats in red, lake state in green and services in blue). Colours of the three-band composite image were scaled with a minimum-maximum scaling method where the lower value of a band equals 0 and the highest value equals 255. All raster layers were done in ArcMap 10.6.1.

## Results

### Determining regional baselines

We observed a considerable amount of overlap suggesting similarities in morphometric and chemical properties among lakes of certain ecozones using linear discriminant analysis (LDA) (Figure S3). Overlap generally occurred between adjacent ecozones and within a common physiographic region (Figure S1). For example, lakes in the Prairies, Boreal Plains, and Taiga Plains ecozones, all from the Interior Plain physiographic region, had similarly high secchi depth, high conductivity, and reduced watershed slope. This suggested that these ecozones could be merged into a large lake region with shared physical and chemical baseline conditions (see Table S1). Based on the amount of overlap observed in the LDA reflecting shared properties among ecozones, and on physiographic spatial proximity, we aggregated several ecozones into five “lake regions” (Figure 2a). These were renamed from West to East as the “Pacific” (*n* = 69), the “Mountains” (*n* = 137), the “Plains” (*n* = 165), the “Continental” (*n* = 220), and the “Atlantic” (*n* = 68) lake regions (Figure 2b). The LDA performed with the new lake regions was more robust, where the two first canonical variates explained 49.19% and 40.30% of the total variance compared to 52% and 29% for the LDA among the original ecozones. Attribution of lakes to their original group was also improved by using the discriminant functions of the new aggregated lake regions. Classification was correct with a mean of 85% using lake region as the grouping when compared to 76% when using ecozones (Table S2). Table 2 describes the variability of the unimpacted reference lake conditions for each lake region.

**Figure 2.**
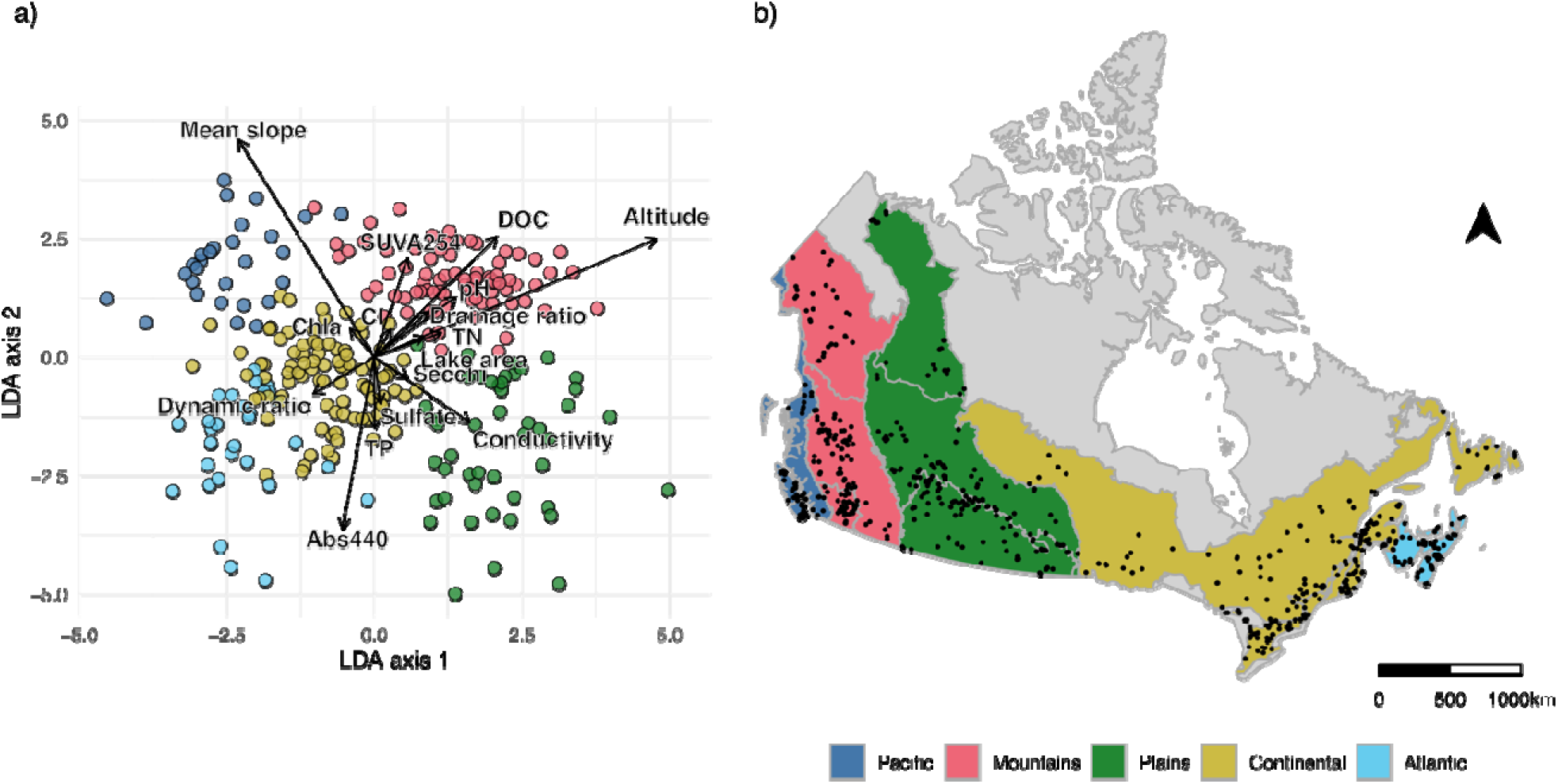
Lake region delineation: a) Linear discriminant analysis (LDA) of low impact lakes using 15 chemical and morphometric variables differentiating lake regions. Points are discriminant scores of sampled low human impact lakes; arrows are the discriminant vectors of chemical and morphometric variables. b) Lake region map with merged ecozones; Pacific (Pacific Maritime ecozone), Mountains (Semi-Arid Plateaus, Boreal Cordillera and Montane Cordillera ecozones), Plains (Prairies, Boreal Plains and Taiga Plains ecozones), Continental (Boreal Shield, Mixedwood Plains and Atlantic Highlands ecozones), and Atlantic (Atlantic Maritime ecozone). Points are the lakes sampled by Lake Pulse network.

**Table 2.** Median values of low human impact lakes variables (not transformed) within the new lake regions.

| **Variables** | **Pacific** | **Mountains** | **Plains** | **Continental** | **Atlantic** |
| --- | --- | --- | --- | --- | --- |
| Secchi (m) | 5.85 | 4.10 | 1.66 | 3.28 | 2.16 |
| TN (mg/L) | 0.09 | 0.20 | 0.54 | 0.15 | 0.14 |
| TP (µg/L) | 6.95 | 15.49 | 32.56 | 12.38 | 15.81 |
| Chl *a* (µg/L) | 0.98 | 2.18 | 4.13 | 2.21 | 2.87 |
| DOC (mg/L) | 1.57 | 7.81 | 21.53 | 6.56 | 6.54 |
| SUVA254 (L/mg-m) | 5.39 | 5.14 | 3.55 | 5.49 | 8.02 |
| A440 (1/m) | 0.24 | 0.85 | 1.38 | 1.27 | 2.70 |
| Chloride (mg/L) | 1.26 | 0.84 | 5.35 | 1.23 | 3.31 |
| Sulfate (mg/L) | 1.99 | 5.92 | 12.48 | 2.39 | 1.26 |
| C:N | 23.48 | 48.45 | 45.22 | 50.23 | 57.27 |
| C:P | 1064.09 | 1141.10 | 1698.44 | 1294.14 | 1131.43 |
| N:P | 31.77 | 20.97 | 33.63 | 25.16 | 18.70 |
| Conductivity (µS/cm) | 43.14 | 158.87 | 387.76 | 113.37 | 36.00 |
| pH | 7.35 | 8.30 | 8.65 | 7.75 | 6.83 |
| Drainage ratio | 26.96 | 24.49 | 12.47 | 15.11 | 12.47 |
| Mean depth (m) | 9.88 | 6.02 | 1.17 | 3.84 | 1.29 |
| Lake area (km<sup>2</sup>) | 1.98 | 1.17 | 0.80 | 1.56 | 0.79 |
| Mean slope | 27.87 | 12.79 | 1.92 | 5.45 | 4.19 |
| Altitude (m) | 215.00 | 850.00 | 476.00 | 240.50 | 95.50 |
| Dynamic ratio | 0.16 | 0.20 | 0.82 | 0.35 | 0.97 |

The LDA discriminant function indicated that altitude, mean watershed slope, colour (Abs440), DOC, and conductivity were the variables that best distinguished lake regions (Figure 2a, Table S3 and S4). We observed that Pacific lakes were generally deep (highest mean depth and low dynamic ratios), unproductive (high secchi depth, low TN, low TP, low Chl *a* and low DOC), and with steep catchment slopes. Mountain lakes were typically quite clear (high secchi), contained little chloride, and were, unsurprisingly, generally located at much higher altitudes. In contrast, Plains lakes were located within shallow catchments, and were the most naturally productive and ion rich lakes. Indeed, baseline values of TN, TP, DOC, and conductivity were twice as high as in other regions, but secchi, and SUVA254 were lowest. Lakes of the Continental region generally exhibited median values falling in the middle of the distribution of most variables. Atlantic lakes tended to be small and shallow (second lowest mean depth and high dynamic ratio), located at low altitudes, and rather coloured (high Abs440) with low conductivity. Also, the Atlantic was the only region where pH was acid (< 7) for most lakes. Conditions of reference lakes in some regions, TN in the Plains for example, were higher than those considered high in all other regions. As such, regional reference conditions need to be established rather than a national average when assessing the deviation for lake ecological state or relative lake health (Table 2).

### Deviation of impacted lakes from reference

To characterize the deviation of lakes from reference conditions, we compared how morphometric, chemical, and land use characteristics of highly impacted lakes deviated from the low impact ones (Figure 3). These deviations were calculated using region-specific z-scores, allowing to compare the direction and/or magnitude of changes among lake regions. For morphometry, impacted lakes generally had lower values compared to reference (Figure 3a), indicating that these lakes tended to be shallower, smaller, had shallower watershed slopes, lower drainage and dynamic ratios, and were located at lower altitudes. The Plains region was an exception to this trend as most morphometric variables had positive deviations, and impacted lakes tended to be larger with higher drainage and dynamic ratios, and located at higher altitudes. Other exceptions were observed in the Pacific region (higher dynamic ratio) and in the Atlantic (deeper lakes with a steeper slope). Finally, while all morphometric variables from impacted lakes of the Continental region were similar to reference, there were large differences for some variables in other regions, namely much lower slope in the Pacific, much lower altitude in the Mountains, and much lower dynamic ratio in the Atlantic.

**Figure 3.**
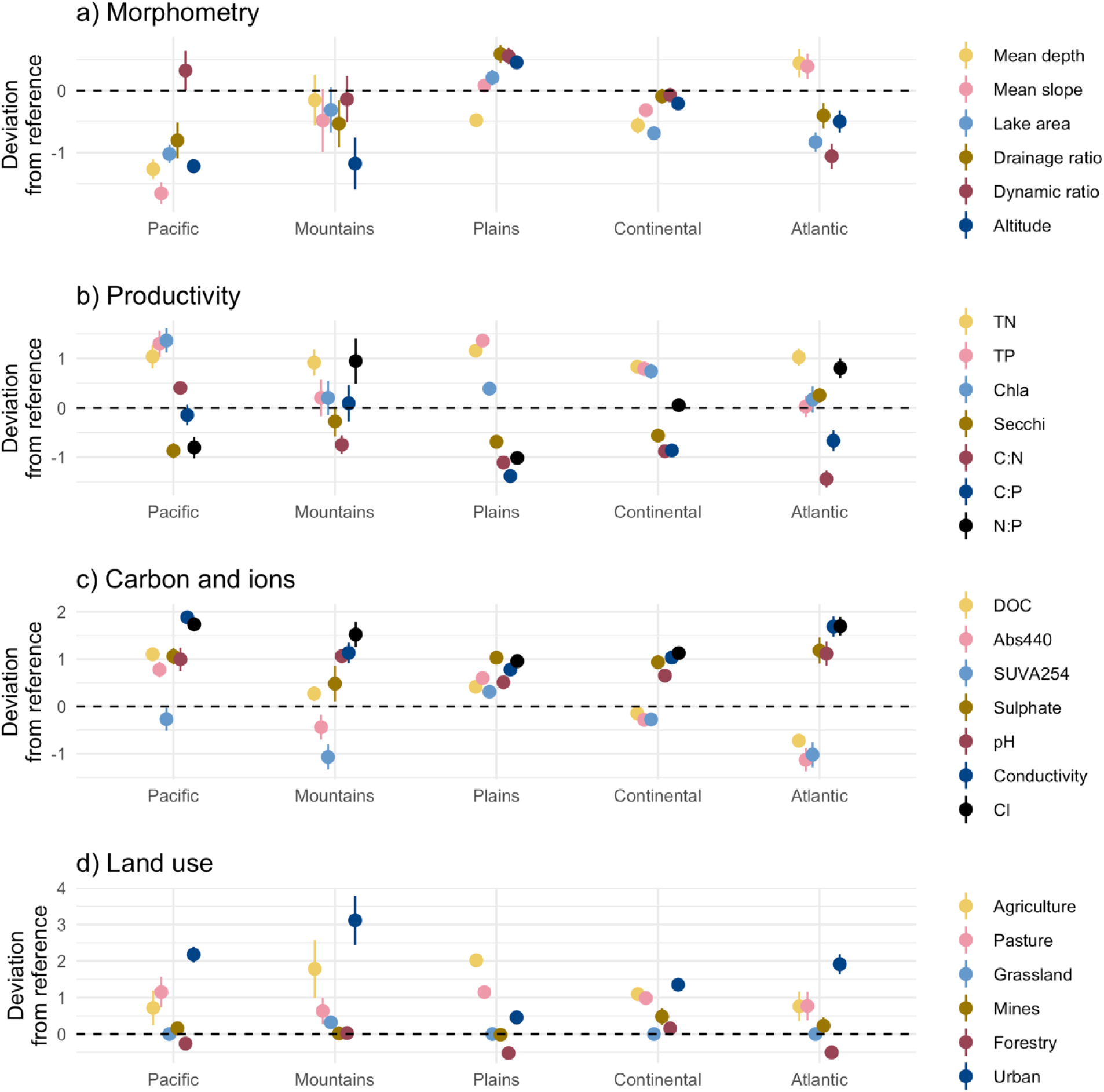
Line plot representing deviation of high impact lakes from reference conditions among different lake regions. Points are the mean values coloured by variable; lines are the standard errors. The dotted horizontal line represents the regional reference. Panel a) represents physical and morphometric features; b) Nutrients, chlorophyll, secchi, and changes in elemental ratios; c) Carbon, spectral properties, and ions, d) Land use in watersheds.

For the deviation of chemical variables, we visualized nutrients, Chl a and secchi separately from variables related to changes in carbon (DOC and spectral properties), pH and ions (Figures 3B and C, respectively). TP and Chl a which are classically used to characterize eutrophication, were higher in impacted lakes for several regions, but not all (Figure 3b). Concentrations in the Mountain and Atlantic regions were similar to reference, and although there was an increase in TP in the Plains, there was no corresponding increase in Chl a. Secchi, which is also classically used to assess eutrophication, was lower in impacted lakes in all regions except in the Atlantic where it was on average higher, and as such highly impacted lakes were counterintuitively more transparent. TN, however, increased rather strongly and consistently across all regions. Fluctuations in nutrients were also observable in the relative changes in N:P ratios where regions responded rather differently. When C:N and C:P ratios were considered, the responses were equally divergent. If TN increased in all cases and DOC stayed the same, the ratios should have all decreased. This however was not always the case as DOC increased regionally in impacted lakes in the Pacific region, decreased in the Atlantic, but remained unchanged in the others (Figure 3c). Spectral measures of abs440 and SUVA254, which represents coloured and terrestrial DOC composition respectively, were largely similar between lakes of low and high impact with some notable decreasing trends in both the Atlantic and Mountain Regions and an increasing trend of abs440 in the Pacific. Cl^-^, conductivity, sulfate and pH increased consistently across all regions in impacted lakes, but the magnitude of the response for chloride was the strongest.

Across regions, not surprisingly, differences in land use for highly impacted lakes were driven predominantly by increasing agriculture or urban expansion in their watersheds (Figure 3d). Urban expansion was the main anthropogenic change across all regions except for the Plains where agriculture was most dominant. Mountain and Continental lakes were strongly affected by both drivers, whereas the Atlantic and Pacific were primarily impacted by urban development. We used a PCA to look for correlations among the deviations from reference lakes of different variables across regions (Figure 4). This allowed us to determine more generalized patterns of changes in chemical variables in response to changes in physical and land use features. Axis one separated impacted lakes between those influenced from by urban and agricultural land use change, where those lakes impacted by urban expansion in their watershed tended to be smaller, clearer, with lower nutrients, located at lower altitudes, and associated with higher Cl^-^ concentrations. Those impacted by agriculture and pasture, tended to be shallower, had higher drainage and sediment dynamic ratios, with higher TP, TN, Chl a and DOC concentrations. As Cl^-^ tended to be more indicative of an urban signal and TN, one of agriculture, and both had a consistently strong deviating response across all regions (Figure 3b and c, Figure S4), we used both these variables as our indicator of altered lake ecological state.

**Figure 4.**
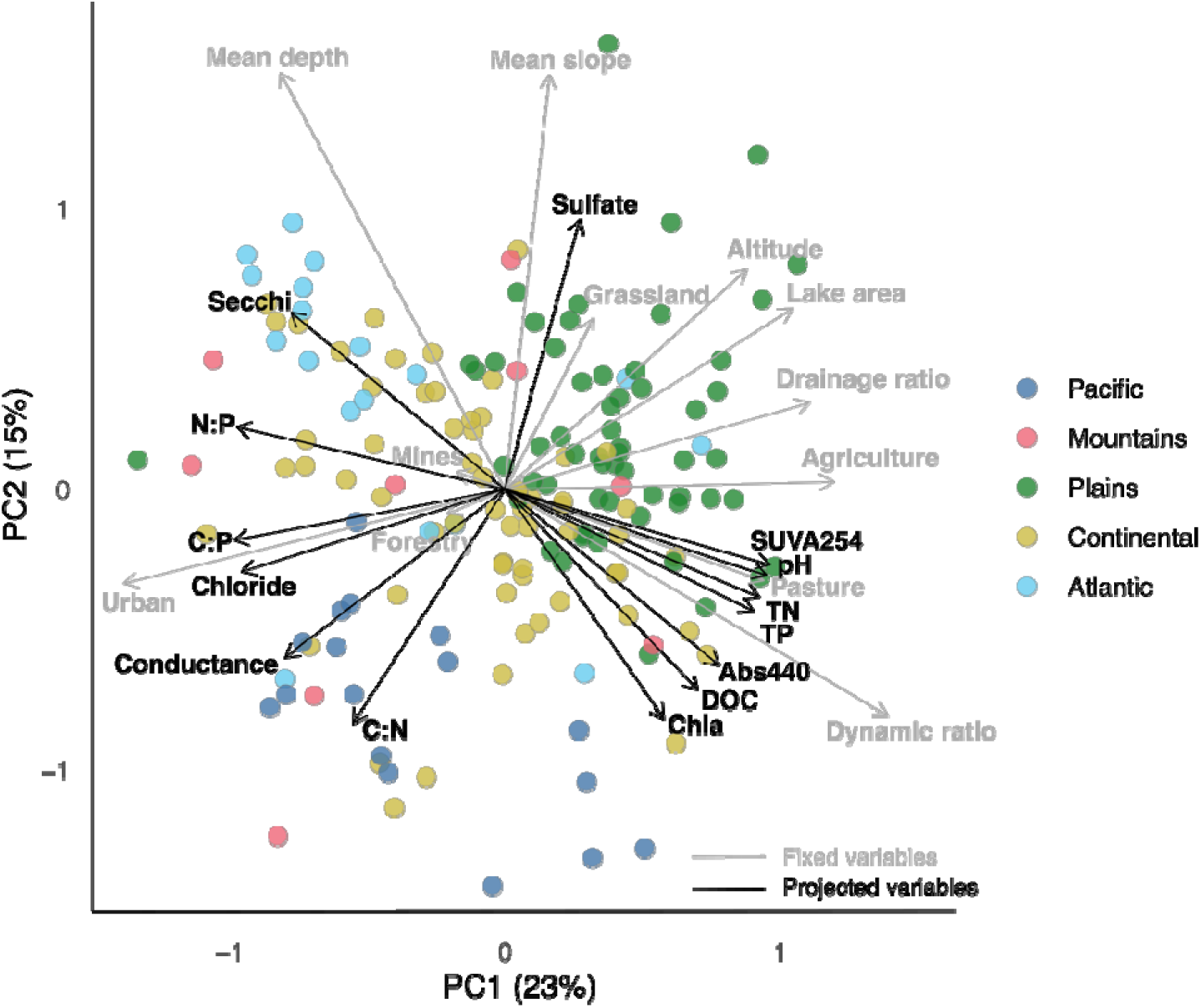
Principal component analysis (PCA) of deviations from reference conditions across passively (fixed) morphometric and land used variables (in grey) and projected chemical variables (in black). Results are depicted in scaling 2 where angles between arrows approximate correlation between variables.

### Social-ecological biogeography: creating gradients across Canada

Using TN and Cl^-^ as indicators of human impact, we created a national map of lake ecological state based on sampled lakes (Figure 5a). Altered lakes were scattered across Canada, but dense clusters were observed in Southern Ontario and across the Prairies. Regional patterns in lake state attributed to Cl^-^, or TN concentrations, or a combination of both emerged (see individual maps of lake state based on chloride or on TN, Figure S5). Newfoundland, Nova Scotia, Northern Ontario, and Vancouver area were predominantly altered due to Cl^-^, while alteration in the central part of the Prairies (Manitoba, Saskatchewan, and Alberta) was attributable to TN. Alteration in the middle of British Columbia, the southern part of the Prairies, Southern Ontario and the Lower St. Lawrence in Quebec was due to a combination of both. The northern regions (Northern British Columbia, Alberta, Saskatchewan, and Quebec) and Western Ontario showed little change in lake state.

**Figure 5.**
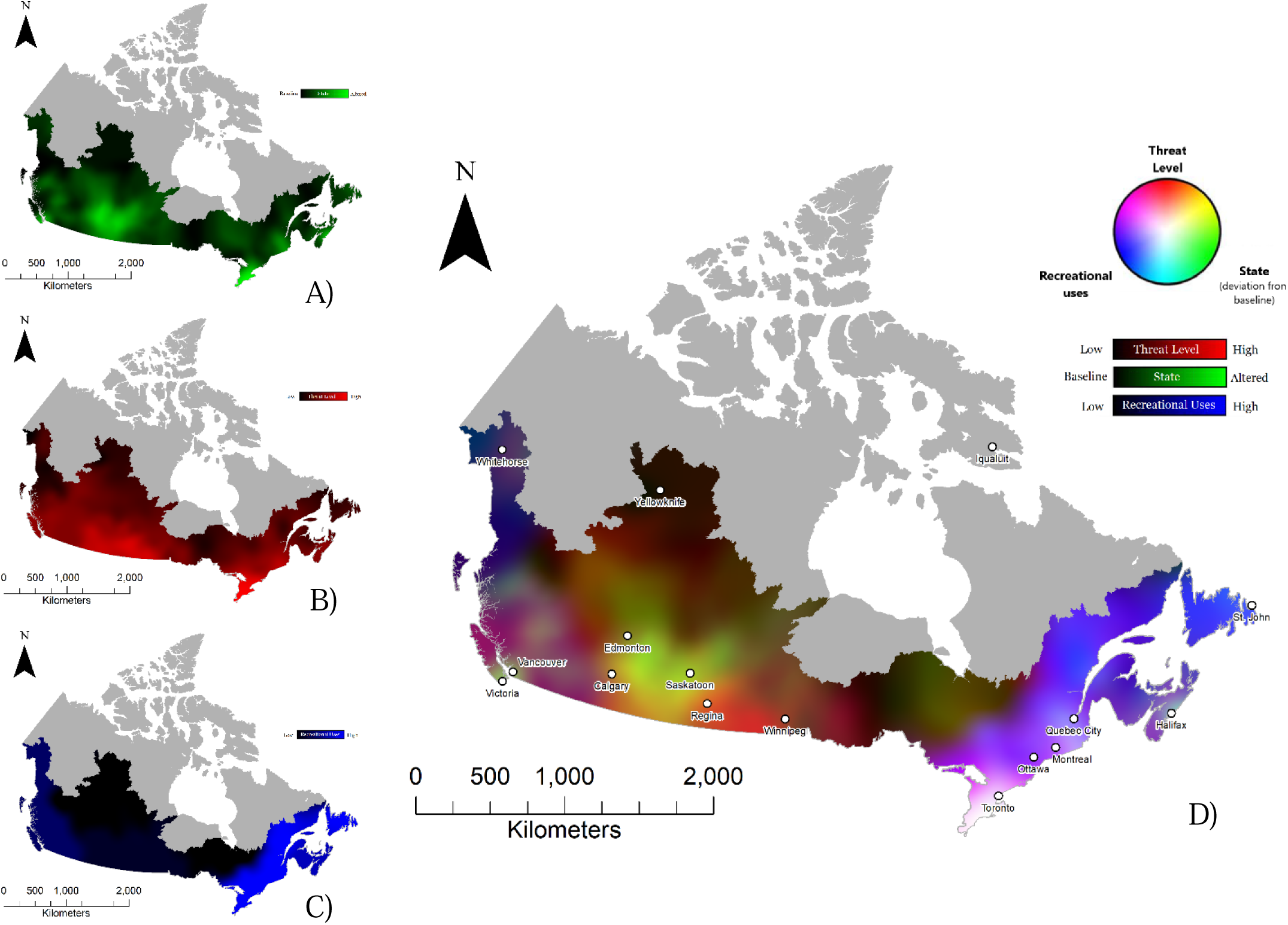
Single coloured maps of ecological state, threat level and recreational use (left) with their combination (right) using red, green, blue (RGB) additive colour scheme. a) Ecological state of the lakes relative to reference conditions (2017-2019); b) Threat levels in the sub-watersheds (2017); c) Recreational uses in the major watersheds (1996, corrected for 2016 population density); d) Composite map created by overlaying the maps on the left.

Threat levels (Figure 5b) were higher in the southern part of Canada, which is coherent with higher population densities. However, northern areas were also subject to threats, mainly climate change and habitat loss. The highest overall threat levels were found around the Great Lakes and in the southern part of the Prairies. In these regions, all of the threats considered were present at a very high level. Exceptions were habitat loss and climate change in Southern Ontario (moderate level) and for invasive species in the Southern Prairies (low level). Regions within the St. Lawrence watershed were the only ones affected by water overuse whereas pollution level was a widespread threat particularly along the border with the United States (WWF-Canada, 2017).

Recreational use (Figure 5c) was highest in Eastern Canada, followed by the West Coast. Participation in the four categories of recreational ES, swimming and beach activities, sport fishing, power boating, and paddle, and sail boating was highest in the St. Lawrence Drainage Area (Quebec, Newfoundland, and Southern Ontario). In the Maritimes Provinces, participation was also very high in all ES categories, except for power boating. Participation in the Pacific Drainage Area (British Columbia and a part of Alberta) was substantial, particularly for sport fishing and swimming, but the frequency of participation (days/km^2^ of watershed) was lower than in Eastern Canada. Use in the Nelson River Drainage Area (Prairies) was low compared to the other three regions, but fishing and swimming were the most popular activities. In the less densely populated northern regions, overall use of recreational ES was low, but sport fishing remained a major ES (Environment Canada, 1999).

On the composite map (Figure 5d), the dominant colour in Eastern Canada changes from purple in the north to magenta in the south, reflecting the high recreational ES (blue) demand in this region with threat level (red) intensifying from north to south. We also note the presence of cyan patches in Southern Quebec and Nova Scotia, particularly around Quebec City and Halifax respectively, implying the presence of altered lakes actively used for recreation. The tip of Southern Ontario appears in white, indicating that lakes were altered, threatened, and intensively used. In Western Canada, similar colour gradients as in the East emerged, but shades tended to be darker. This suggests that the same components (recreational uses and threat level) were present, but at lower levels compared to the East. Threat level also intensifies on a north-south gradient in the west, from Whitehorse to Vancouver. The few diffuse green spots indicate areas where lakes were particularly altered. In Southern Manitoba, the red to orange colour suggests that despite the strong presence of threats, the lakes in this region have suffered minimal alteration compared to their reference state. In Southern Saskatchewan, however, the yellowish colour indicates significant levels of threat and lake alteration. In the middle of these two provinces, there is a light green clump signifying the alteration of several lakes, but with a lower threat level than those located to the south. Recreational use does not emerge in these regions as the level of participation was considered low compared to the rest of Canada. The main colour obtained in Northern Canada (Northern Quebec and Ontario, Northern Saskatchewan and Alberta and Northwest Territories) was dark brown, indicating a low level of threat, little to no recreational use, and low levels of lake alteration.

## Discussion

In this study, we determined the regionally specific reference conditions of lake ecological state for five lake regions across Canada. Reference conditions were based on shared physical and chemical characteristics of unimpacted lakes from ecozones with the same physiographic properties. With this information, we were able to evaluate how severely lakes with greater human activities in their watershed deviated from reference conditions, and found two consistent indicators of human altered lake ecological state across Canada: an increase in total nitrogen (TN) and chloride (Cl^-^) concentrations. The ecological state of several lakes determined using these indicators were then geographically linked to two other aspects of overall lake state, which included a lake’s vulnerability to future threats, and recreational ecosystem service (ES) use. By combining lake ecological state to aquatic ecosystems threats, and ES demand using an additive colour scheme, we created the first national social-ecological geography of Canadian lakes. This novel approach allows us to identify areas where lakes are most vulnerable to being altered or lose their capacity to provide ES, that can serve as priority sites for conservation or restoration to sustain or improve water quality.

### Determining lake ecological state: regional baselines and human indicators of alteration

Understanding how severely a lake is altered by human activities in its watershed requires the knowledge of reference conditions prior to those changes. Spatial differences in the underlying geology that influence the topography, morphometry and land cover will likely create regionally distinct lake baseline types (Boyce, 2015; Read et al., 2015), which are essential to characterize when carrying out comparisons at the continental scale (Cheruvelil et al., 2013). The stratified design of Lake Pulse sampling that included a series of low or unimpacted lakes across most of the country allowed us to determine these reference conditions (Huot et al., 2019). We found that unimpacted lakes from several Canadian ecozones adjacent to one another with shared physiography had similar morphometric and chemical baseline traits. Landscape and watershed characteristics such as elevation and mean slope were the most important variables in discriminating lake properties between regions, followed by conductivity, DOC, and to a lesser extent nutrients. This finding highlights the role of physiography in creating the lake regions across Canada (Boyce, 2015), suggesting that climate and vegetation type related to ecozones had less of an overall influence on reference lake physical chemical characteristics.

The deviation of highly impacted lakes from their regional reference allowed us to evaluate which chemical variables were most consistent in capturing a lake’s response to anthropogenic changes in its watershed. Typically land use change results in higher nutrient inputs with subsequent increased algal biomass and loss of transparency in lakes (Carpenter et al., 1998; Nielsen et al., 2012). Most regions behaved as expected and experienced cooccurring higher concentrations of TN, TP, and Chl *a*, with reduced secchi due to increased turbidity (Minns et al., 2008; Webster et al., 2008). However, higher TN in impacted lakes was more consistently observed across the lake regions, and not always accompanied by higher TP and Chl *a* or transparency loss. Increases in TP, and as a consequence algal biomass, may have been dampened in regions where urban expansion was dominant due to legislative reductions from point source sewage inputs or well managed septic systems (Budd & Jones, 1978; Schindler, 2006). One exception was in the naturally P rich region of the Prairies (Table 2) where despite a significant increase in both nutrients, the increase in Chl *a* was not as marked. This could be explained by a threshold response of Chl *a* to increasing TN (Filstrup et al., 2018). Another possible explanation of why TN was more indicative of human impact in the watershed is the relatively higher applications of N as compared to P fertilizers on agricultural lands (Glibert et al., 2014; Goyette et al., 2016). Excessive N inputs relative to P, in conditions when both are high, is cause for concern as higher N:P ratios tend to favor the establishment of more toxic species of algae and cyanobacteria which impinges on multiple valuable aquatic ES (Glibert et al., 2014; Leavitt et al., 2006; Scott & McCarthy, 2010; Taranu et al., 2017). TN has been shown to increase even with modest human activities in the watershed (Tremblay et al., 2020), and emerged as early warming human impact indicator more so than either TP or Chl*a* in this study. TN was a consequence of both urban and agricultural expansion, but was the main agricultural indicator.

Another striking and consistent difference between human impacted and reference lakes among regions was higher Cl^-^ concentrations. In northern freshwaters, this increase in Cl^-^ concentrations is primarily attributed to road salt used for de-icing during winter, which explains why it was most associated with urbanization in the watershed (Canadian Council of Ministers of the Environment, 2011; Chapra et al., 2009; Dugan et al., 2017). Regions with the highest road salt use in Canada are Southern Ontario, Quebec, and the Maritimes (Environment Canada & Health Canada, 2001; Evans & Frick, 2001) and salt application is also reported to be high in the Prairies (Morin et al. 2000), consistent with our observations of altered Cl-concentrations in these regions (Figure S5a). The environmental impact of increased Cl^-^ concentration in lakes is of growing concern particularly with changing climate conditions where warmer average winter temperatures and changes in precipitation may require more salt for de-icing (Kaushal et al., 2021). High levels of Cl^-^ can have adverse effects on aquatic life with chronic toxicity occurring at 250 mg/L (Evans & Frick, 2001), with recent experimental work showing that adverse effects and shifts in community composition at even lower concentrations of Cl^-^ highlighting the need to downscale toxicity levels even further (Hébert et al. 2022; Hintz et al. 2022). Increased salinization may also impact physical lake properties, including stronger and more prolonged stratification, with cascading repercussions on major biogeochemical cycles (Dugan et al., 2017; Kaushal et al., 2005). As such increased Cl^-^ concentrations may influence the provisioning of several ES provisioning including sport fishing. However, the overall consequences of increased salinization on lake ecosystem functioning and services remain underexplored. Regardless, Cl^-^ was the strongest human indicator of urban expansion. Both TN and Cl^-^ to water can render drinking water unpotable due to concerns around human health and taste respectively (Howard & Maier, 2007; Karimi et al., 2020; Kaushal et al., 2005) making their combined deviation a good proxy for assessing altered lake ecological state, and vulnerability to provide different ES.

Beyond the clear patterns of TN and Cl^-^ increase in impacted lakes, we also found some interesting anomalies in the deviations from certain chemical variables in some regions. For example, impacted lakes of the Atlantic and Mountains regions were clearer than reference lakes on average, as secchi depth was either slightly higher or unchanged. As Chl *a* concentrations in impacted lakes were similar to reference lakes, this enhanced water clarity could be explained by the lower coloured DOC (abs440, Brezonik et al., 2019). A possible influence for this loss in colour is photobleaching of CDOM which can increase water transparency (Del Vecchio & Blough, 2002; Kalff, 2002). Overall lakes from the Mountain region are on average deeper and at higher altitudes (Table 2) whereas impacted Atlantic lakes, morphometrically, tended to be deeper with a lower sediment-dynamic ratio (Figure 3). For both, this would result in longer water residence times, favoring photobleaching. Indeed, in terms of morphometric, topographic, and landscape variables, most impacted lakes tended to be near or below reference lakes, (with the exception of the Plains), but each region has its peculiarities. One overarching tendency is that impacted lakes across multiple regions were more accessible to human populations. This includes impacted Pacific, Mountains, and Atlantic lakes which were at lower altitudes. In the case of the Pacific region, impacted lakes had much shallower watershed slopes than unimpacted ones, and are presumably easier to develop on. In the Plains, Pacific and Continental regions, impacted lakes were generally smaller and shallower, making them more vulnerable to changes in the landscape due to higher internal loading and reduced volume (Nõges, 2009). Impacted lakes in the Plains had both higher drainage and dynamic ratios resulting in a higher proportion of terrestrial inputs (Kalff, 2002) and a greater proportion of the water in contact with sediments (Håkanson, 1982), respectively. These physical features combined could explain the slightly higher colour in the impacted lakes of this specific region, as their physical properties make them inherently more vulnerable to human pressure.

### A first regional social ecological geography of overall lake state

In order to create the first social-ecological geography of lakes in Canada, we developed an innovative framework that spatially links regional lake ecological state, emerging lake threats and recreational ES use. By colour-coding ecological and social gradients, the RGB composite map (Figure 5) allowed us to visualize hotspots of aquatic ecosystem vulnerability and ES conservation. This approach clearly identified areas where lakes were in demand by people and where there was a risk of losing recreational ES due to threats or lake degradation. From the perspective of preserving recreational ES where there is demand, the colours that interested us were white (high alteration, high threats, and high recreational use), cyan (high alteration and high recreational use), and magenta (high threats and high recreational use). These colour patterns do indicate locations where demand for services can be expected to coincide with lake degradation and/or threat, and thus should be monitored as a precaution, although they cannot definitively determine whether or not recreational services will be lost over time.

Through this framework, we found that populated urbanized areas were of greatest concern. The Southern tip of Ontario, which includes the Greater Toronto Area and accounts for nearly 20% of the country’s population (Canada Population, 2021), appeared in white on the map due to a combination of lakes being intensively altered, threatened, and used. Other areas of concern include the Lower St. Lawrence River between the cities of Montreal and Quebec, and the Halifax region in Nova Scotia, which all exhibit a colour approaching light cyan, indicating high recreational use combined with altered lake ecological state and a moderate threat level. All these urban locations should be monitored as degraded lake state could potentially lead to the loss of lake services. The same colour combinations were found in British Columbia around the cities of Vancouver et Victoria area, but according to the data we had, recreational use on the West Coast was lower, resulting colour tends to be a lighter green with altered lake ecological emerging as a more prominent component overall. Areas coloured magenta were found in Southeastern Ontario, Southern Quebec and in the lower part of British Columbia. Here recreational use and threat level were high but lakes on average are not yet significantly degraded. Finally, in central Canada, particularly in the Saskatoon, Saskatchewan and Edmonton, Alberta areas, lakes were altered and threatened, and although demand for recreational services is low, lakes may be at significant risk, primarily from agriculture, but also urbanization.

### Limits and considerations

One limitation to our study, is that our indicators of altered lake ecological state, TN and Cl^-^ do not necessarily impinge directly on the delivery of the four recreational services assessed here, but may act as a proxy for some. For example, Health Canada’s Guidelines for Canadian Recreational Water Quality (2012) related to swimming has threshold values for cyanobacteria and *E.coli* related to the risk to human health in case of direct contact to skin. High concentrations of TN are known to promote toxic algal and cyanobacterial blooms (Glibert et al. 2014; Monchamp et al. 2014; Zaranu et al. 2017), and indirectly linked to loss of this ecosystem service. Similarly, Cl^-^ may through food web effects (Dugan et al. 2017; Hintz et al. 2022), influence sport fisheries although that remains underexplored. A more appropriate ecosystem service to assess with nitrate (as part of TN) and Cl^-^ is the loss of drinking water as increasing concentrations of both can render water unpotable due to human health concerns and taste respectively (Howard & Maier, 2007; Karimi et al., 2020; Kaushal et al., 2005), but we did not find easily accessible data around that ecosystem service across Canada.

There were also several limits with the recreational ES dataset we used. First it was dated from 1996, and although adjusted for the current population, it is possible that recreational habits have changed over time and no longer accurately reflect current demand. Second, the ES dataset covered broad watershed areas across the country, and was not as spatially explicit as other components in our model. The recreational ES data were also only available in days of use per square kilometer of major watershed, which reflects the intensity of use over a given area, not the intensity of use by individuals. Less densely populated areas are disadvantaged as use could be diluted by watershed size, where a sparsely populated area in which individuals use ES intensively can be equivalent to a densely populated area where individual use is infrequent. Furthermore, many of these sparsely populated areas are inhabited by First Peoples who were not considered in the census; given their intimate relationship with natural resources, we argue the need to prioritize the protection of ecosystem services for those Nations who rely directly on them for provisioning (Berkes, 1990; Islam & Berkes, 2016).

## Conclusion

Our holistic social-ecological approach allowed us to synthesize a large amount of information about overall state of lakes across Canada. We have shown that there are broad regions where restoration and conservation efforts should be focused given the combined influence of alteration of lake ecological state, emerging threats, and ES demand. Despite the caveats related to some of the data used in our study, there is strength in quantifying ES directly to assess how they are impacted and threatened using our integrated approach. Future work should try to do this more explicitly as although ES are easily identified and defined (MEA 2005), they are often not quantified (Raudsepp-Hearne et al. 2010). Lake ecological state also needs to be put into regional context related to reference conditions, and indicators of ES provisioning as well as lake state may need to be revisited to allow for more commonly measured proxies to be used in broader spatial and historical evaluations (St-Gelais et al. 2020). Our approach is both powerful in its easily understood visual representation, and adaptable at different spatial scales. We recommend its use at finer regional scales to identify hotspots relevant to management. Furthermore, we encourage that it be adapted to ES of regional concern in relation to metrics or proxies known to differently impact and threatened lake ES at more local scales. In a context of climate change adaptation, the demand for freshwater recreational ES as well as others will increase. It is therefore essential to secure the ES lakes provide by reducing pressures and anticipating changes in state. Our study offers a useful and novel tool for management to evaluate ES vulnerability, helping identify places to intervene for long-term ES sustainability.

## Acknowledgements

We would like to thank all of the participant of the Lake Pulse Strategic Network from the collection and processing of samples, with a special thanks to lead PI Y. Huot (University of Sherbrooke) and all members of the coordinating, sampling, and analytical teams. We thank A. Prince for help with all map figures, including the RGB model, and R. Siron from OURANOS for discussions. Funding for this study came from Lake Pulse NSERC Strategic Network, a MITACS with OURANOS, and NSERC CRC in Aquatic Ecosystem Science and Sustainability to RM as well as a BIOS² CREATE program student scholarship to AD. This work is a contribution to the Groupe de Recherche Interuniversitaire en Limnologie (GRIL), an FQRNT Strategic cluster. We acknowledge that this work was carried out on lands belonging to many First Peoples across the mid northern areas of Turtle Island, and share the desire to restore and protect them now and for future generations.

