## supplementary material for "A social-ecological geography of southern Canadian Lakes"

**Text S1**

*Study area*

Canada is a northern country that stretches from the Pacific to the Atlantic Ocean across a wide variety of natural landscapes. In terms of physiography (Figure S1 a), starting in the western part of the country, there are two major mountain ranges known as the Coast Mountains and the Rocky Mountains, surrounding the Okanagan Valley that make up the Canadian Cordillera (Acton et al., 2015). Next, from the province of Alberta to the eastern end of Manitoba are the Interior Plains, characterized by generally flat terrain and rich soils (Wiken, 1996). Then, the Canadian Shield, the largest physiographic region in Canada, extends from northern Saskatchewan to Newfoundland where there are vast forests, numerous lakes and rivers, and bedrock exposures (Renwick, 2009). The southern part of Ontario and Quebec make up the physiographic region known as the Great Lakes and St. Lawrence Lowlands, which is predominantly a plain that stretches along the St. Lawrence River, dotted with hills (Elson, 2010). Finally, at the eastern end of the country, the Maritime Provinces and part of Québec lie in the Appalachian Region, a mountainous region comprising the Appalachians mountain range and an indented coastline (Bone, 2003). In the northern regions of the country, there are the Arctic and Subarctic Regions and the Hudson Bay Lowlands, but these regions were not part of the study area. The physiographic regions are subdivided into ecozones, as shown in Figure S1a.

Canada has a population of 39 million inhabitants that is highly concentrated geographically, with two third of people living within 100 kilometers of the southern border with the United States (Statistics Canada, 2017), and a large proportion of the population concentrated in a few large cities (Figure S1B). Agricultural areas are found mainly in the Prairies, along the St. Lawrence River up to the Great Lakes and on Prince Edward Island (Karimi et al., 2020), as these regions are home to fertile land (Bone, 2003; Wiken, 1996). Most of Canadian lakes were formed by the retreat of the glaciers and surface deposits are generally thin except in the Plains region (Boyce, 2015).

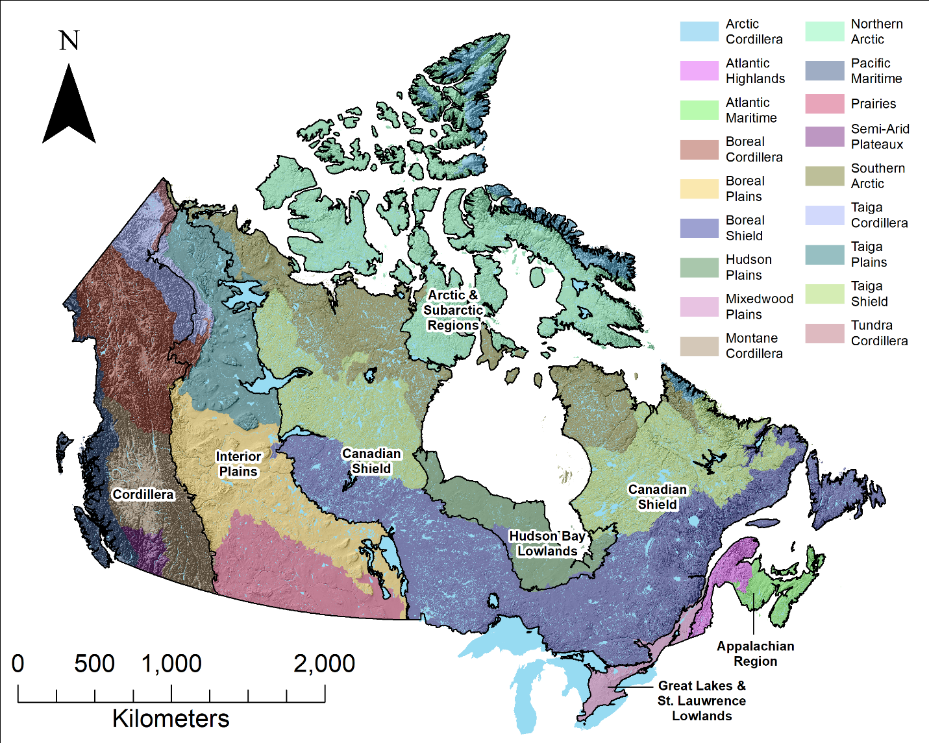

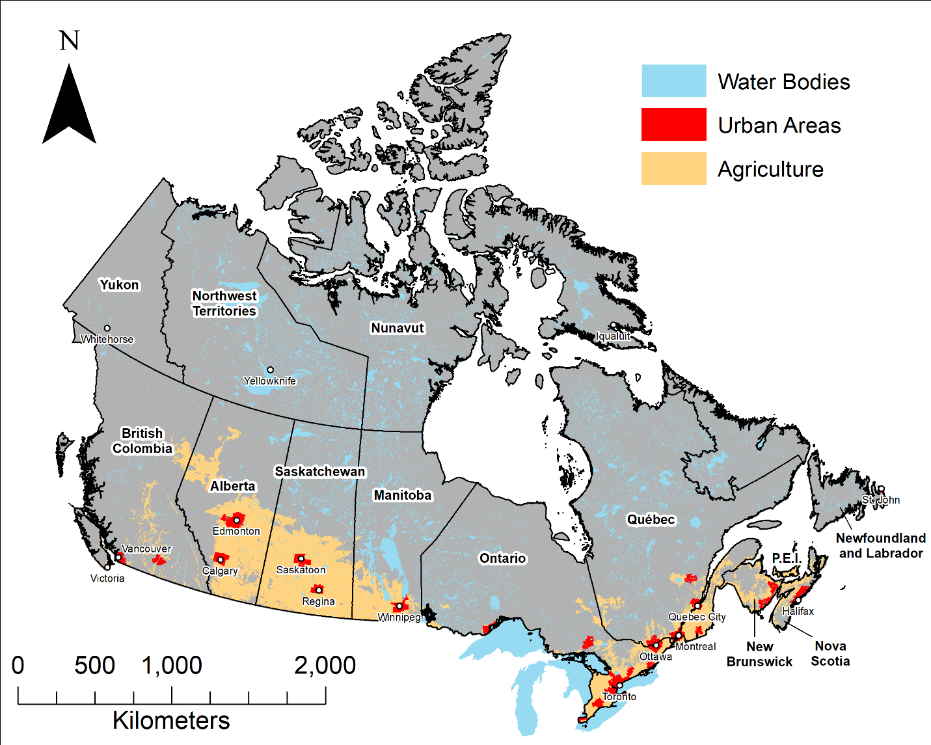

A)

B)

**Figure S1**: Maps of Canada. **A)** Physiographic regions, ecozones and topography of Canada. The physiographic regions are delineated by black lines and their names are shown in black on white on the map; the ecozones boundaries are displayed by the different colors. Source : Physiographic Regions (Natural Resources Canada, 2020), ecozones from the Canadian Council on Ecological Areas (CCEA, 2014), topography from Canadian Digital Elevation Model (Natural Resources Canada, 2017). **B)** Map of Canada's urban (red) and agricultural (yellow) areas and water bodies (blue). Provinces and territories are delineated by black lines with their names in black on white on the map. Major cities and capitals are indicated by white dots. Source: Water bodies from CanVec Series (Natural Resources Canada, 2019), agricultural areas form Landsat LandCover (CEC, 2010) and urban areas from the 2016 Census metropolitan areas and census agglomerations (Statistics Canada, 2016).

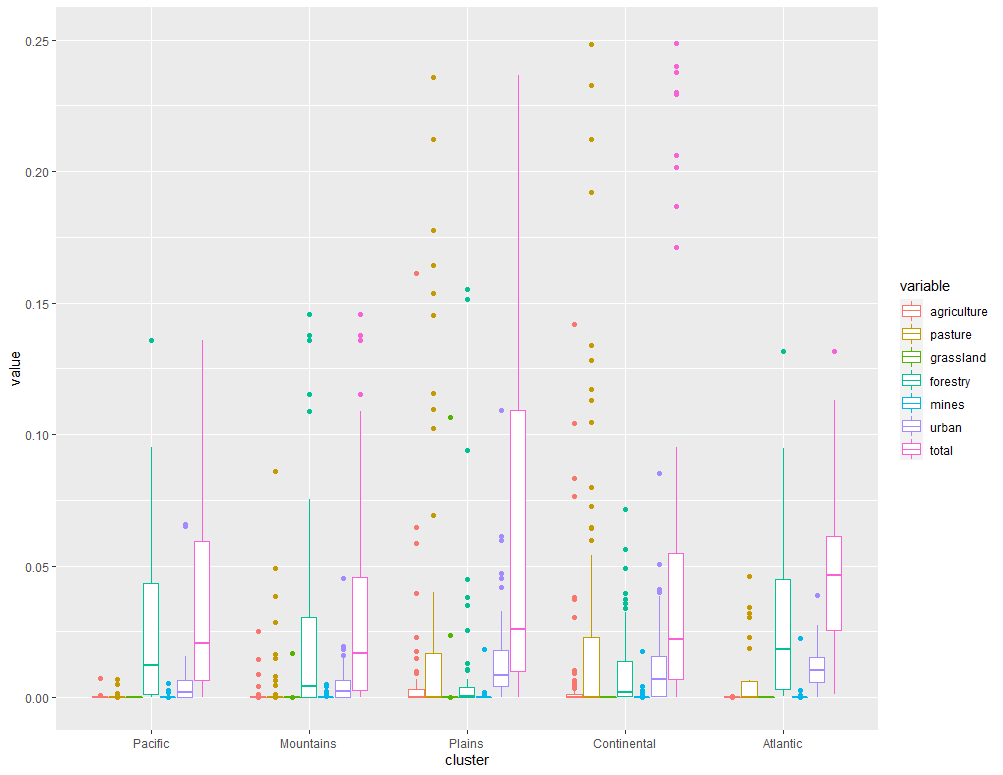

**Figure S2**: Proportion of land use and land cover in the low human impact lakes watersheds by lake region.

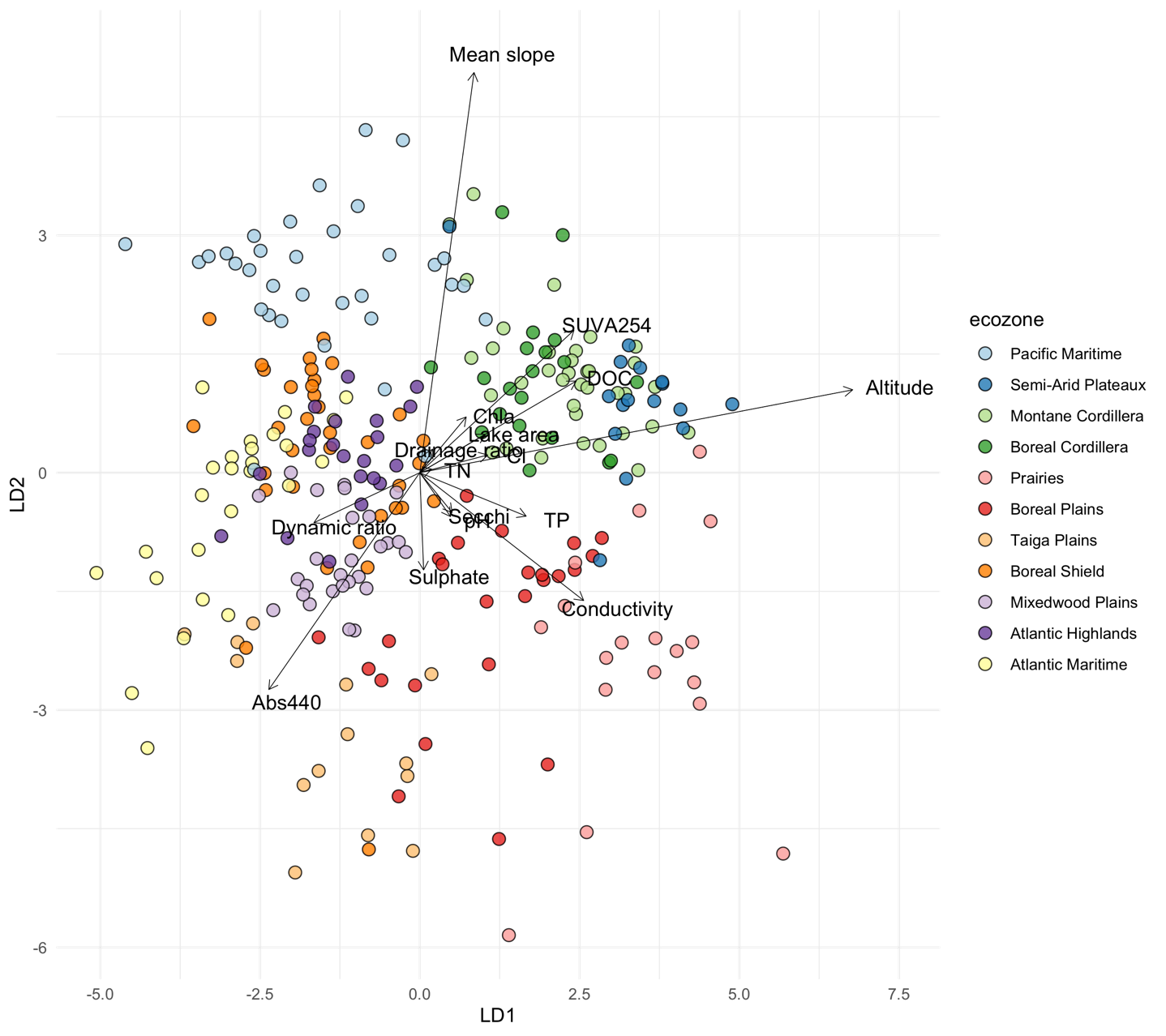
**Figure S3**: Linear discriminant analysis (LDA) using 15 discriminant chemical and morphometric parameters with ecozone as the grouping variable. Points are the discriminant scores of low human impact sampled lakes; arrows are the discriminant vectors of chemical and morphometric variables.

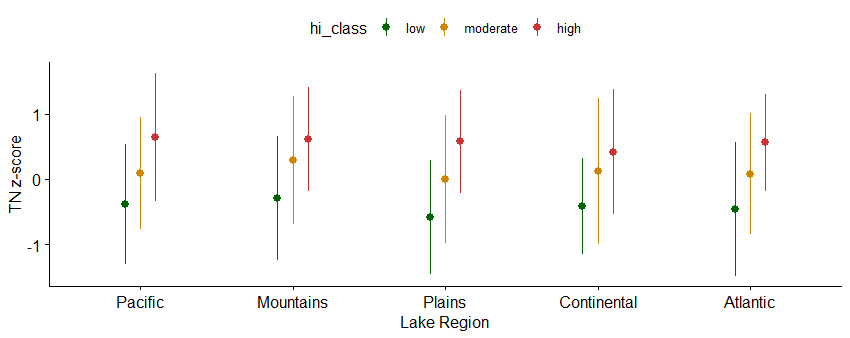

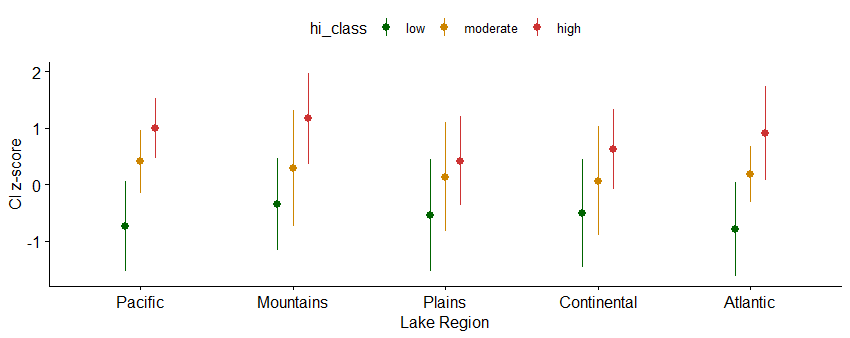

A)

B)

**Figure S4:** Standard scores of A) TN and B) Cl^-^ in the three human impact classes by lake region. Points are the mean values; lines are the standard deviation.

**Figure 5:** Ecological state of the lakes relative to reference conditions (2017-2019); **A)** Deviation in Cl^-1^ concentration; **B)** Deviation in TN concentration

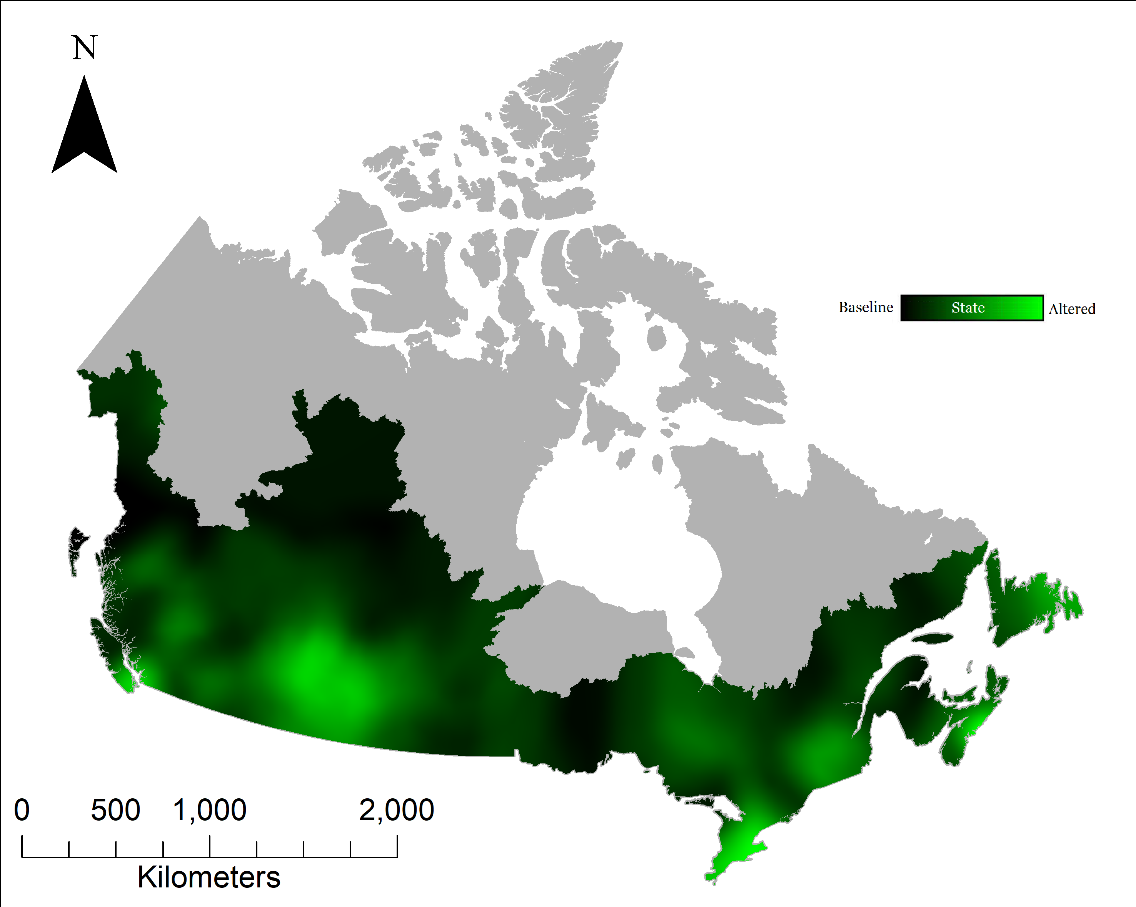

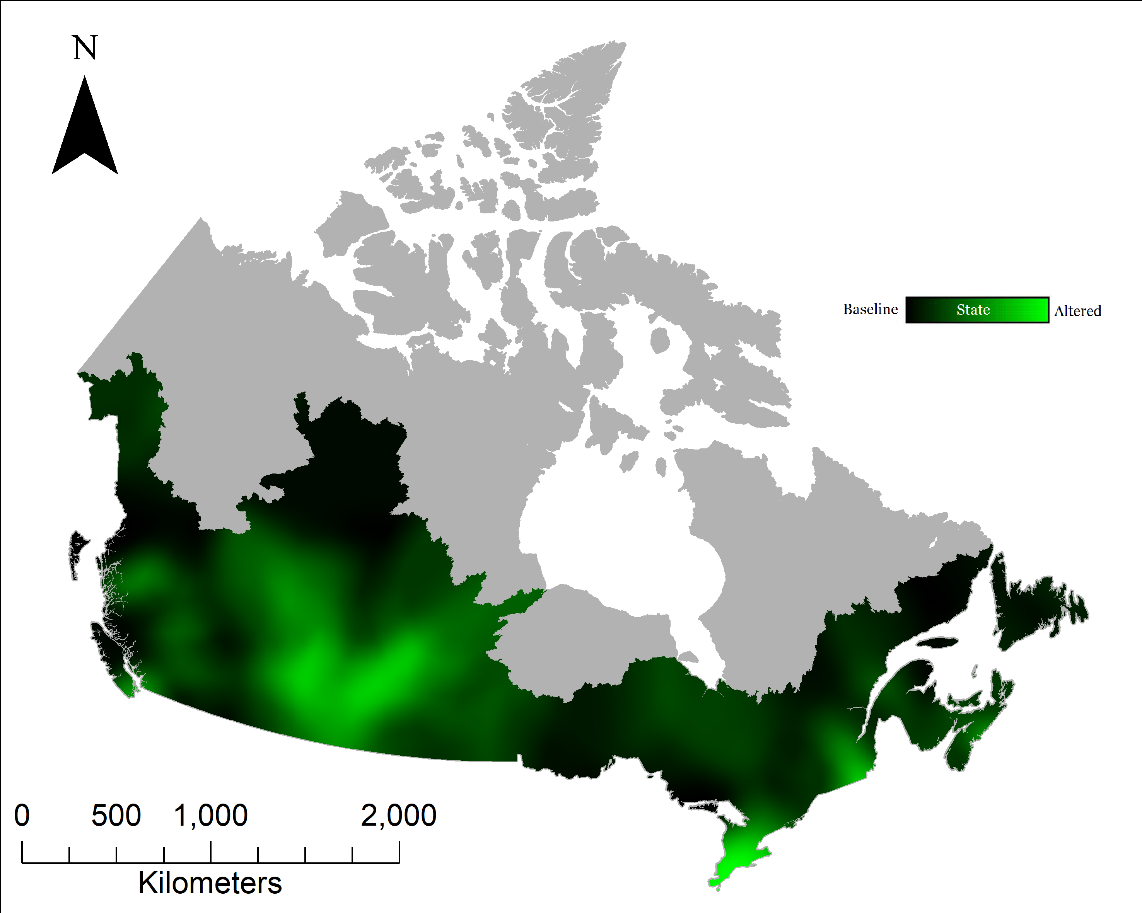

A)

B)

Table S1.
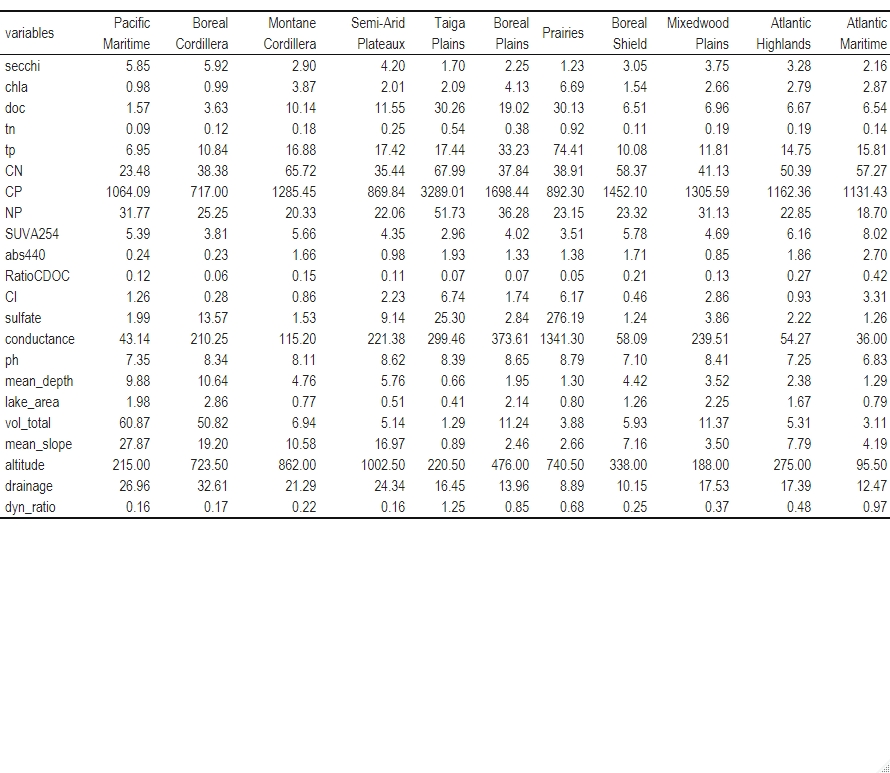
Comparison of mean values for low impact lakes in each ecozones.
Table S2. Comparison of classification table with ecozone or lake region as grouping variable.

| Groups | | Proportion of correct classification | |
| --- | --- | --- | --- |
| Ecozone | Lake Region | Ecozone | Lake Region |
| Pacific Maritime | Pacific | 76 | 73 |
| Semi-Arid Plateaux | Mountains | 63 | 93 |
| Boreal Cordillera |  | 75 |  |
| Montane Cordillera |  | 86 |  |
| Prairies | Plains | 78 | 81 |
| Boreal Plains |  | 80 |  |
| Taiga Plains |  | 86 |  |
| Boreal Shield | Continental | 60 | 89 |
| Mixedwood Plains |  | 96 |  |
| Atlantic Highlands |  | 50 |  |
| Atlantic Maritime | Atlantic | 85 | 88 |

Table S3. Coefficients of linear discriminants with ecozone as grouping variable.

| Variables | LD1 | LD2 | LD3 |
| --- | --- | --- | --- |
| Conductivity | 0.649 | -0.596 | -0.466 |
| TP | 0.417 | -0.162 | -1.054 |
| TN | 0.046 | 0.015 | 0.601 |
| Secchi | 0.126 | -0.115 | -0.022 |
| Chl *a* | 0.181 | 0.188 | -0.009 |
| DOC | 0.610 | 0.357 | 0.190 |
| A_cdom(440) | -0.589 | -0.775 | -0.315 |
| SUVA254 | 0.602 | 0.507 | -0.451 |
| pH | 0.118 | -0.104 | 0.939 |
| Sulfate | 0.015 | -0.307 | 0.040 |
| Chloride | 0.268 | 0.013 | -0.152 |
| Mean slope | 0.221 | 1.185 | -0.046 |
| Lake area | 0.259 | 0.060 | 0.066 |
| Drainge ratio | 0.051 | 0.057 | 0.493 |
| Dynamic ratio | -0.416 | -0.131 | 0.054 |
| Altitude | 1.694 | 0.264 | -0.395 |

Table S4. Coefficients of linear discriminants with lake region as grouping variable.

| Variables | LD1 | LD2 | LD3 |
| --- | --- | --- | --- |
| Conductivity | 0.417 | 0.538 | -0.879 |
| TP | 0.003 | 0.443 | -0.409 |
| TN | 0.215 | -0.113 | 0.964 |
| Secchi | 0.140 | 0.111 | -0.077 |
| Chl *a* | -0.102 | -0.182 | -0.057 |
| DOC | 0.529 | -0.657 | 1.135 |
| Abs440 | -0.148 | 0.938 | -1.596 |
| SUVA254 | 0.147 | -0.558 | 0.169 |
| pH | 0.348 | -0.365 | 0.306 |
| Sulfate | 0.028 | 0.246 | 0.106 |
| Chloride | 0.075 | -0.120 | 0.210 |
| Mean slope | -0.545 | -1.064 | 0.045 |
| Lake area | 0.292 | -0.089 | -0.046 |
| Drainge ratio | 0.233 | -0.233 | 0.260 |
| Dynamic ratio | -0.264 | 0.178 | 0.018 |
| Altitude | 1.203 | -0.633 | -0.498 |

To determine whether chemical and morphometric descriptors could discriminate among the new groups, a second LDA with lakes grouped by lake region (Figure 2 a) was carried out on the same explanatory variables as for the first LDA. The two first canonical variates of the LDA accounted for 49.19% and 40.30% of the total variance respectively. The coefficients of the linear discriminants (Table S2) indicate that the variables that best explain the variation between groups are the same as in the first LDA. According to Figure 2a, the lake regions are distributed along two gradients. The first one is dominated by altitude (with Mountains at high altitude, Atlantic at low altitude and Continental in between), while the second gradient is dominated by the mean slope (with Pacific on the steep slope side, Plains on the low slope side and Continental in between). The coefficients of linear discriminants and the classification table are presented in the supplements (Table S2 and Table S3).
